# Transcriptomics-driven characterisation of novel T7-like temperate *Pseudomonas* phage LUZ100

**DOI:** 10.1101/2022.08.29.505779

**Authors:** Leena Putzeys, Jorien Poppeliers, Maarten Boon, Cédric Lood, Marta Vallino, Rob Lavigne

**Affiliations:** Laboratory of Gene Technology, Department of Biosystems, KU Leuven. Kasteelpark Arenberg 21, 3001, Leuven, Belgium; Institute for Sustainable Plant Protection, CNR. Strada delle Cacce 73, 10135, Torino, Italy

## Abstract

The *Autographiviridae* is a diverse yet distinct family of bacterial viruses marked by a strictly lytic lifestyle and a generally conserved genome organization. We here characterise *Pseudomonas aeruginosa* phage LUZ100, a distant relative of type phage T7. LUZ100 is a podovirus with a limited host range and identified LPS as the likely phage receptor. Interestingly, infection dynamics of LUZ100 indicated moderate adsorption rates and low virulence, hinting towards temperate behavior. This hypothesis was supported by genomic analysis, which showed that LUZ100 shares the conventional T7-like genome organization, yet encodes key genes associated with a temperate lifestyle. To unravel the peculiar characteristics of LUZ100, ONT-cappable-seq transcriptomics analysis was performed. This data generated a bird’s-eye view of the LUZ100 transcriptome and enabled the discovery of key regulatory elements, antisense RNA, and transcriptional unit structures. The transcriptional map of LUZ100 also allowed us to identify new RNAP-promoter pairs that can form the basis for biotechnological parts and tools for new synthetic transcription regulation circuitry. The ONT-cappable-seq data revealed that the LUZ100 integrase and a MarR-like regulator (proposed to be involved in the lytic/lysogeny decision), are actively co-transcribed in an operon. In addition, the presence of a phage-specific promoter transcribing the phage-encoded RNA polymerase, raises questions on the regulation of this polymerase, and suggests it is interwoven with the MarR-based regulation. This transcriptomics-driven characterisation of LUZ100 supports the increasing evidence that T7-like phages should not straightforwardly be marked as having a strictly lytic lifecycle.

**Importance:** Bacteriophage T7, considered the ‘model phage’ of the *Autographiviridae* family, is marked by a strictly lytic lifecycle and conserved genome organisation. Recently, novel phages of this clade are emerging and showing characteristics associated to a lysogenic lifecycle. Screening for temperate behaviour is of outmost importance in fields like phage therapy, where strictly lytic phages are generally required for therapeutic applications. In this study, we’ve used an omics-driven approach to characterise the T7-like *Pseudomonas aeruginosa* phage LUZ100. These results led to the identification of actively transcribed lysogeny-associated genes in the phage genome, pointing out that temperate T7-like phages are emerging more frequent than initially thought. In short, the combination of genomics and transcriptomics allowed us to obtain a better understanding of the biology of non-model *Autographiviridae* phages, which can be used to optimize the implementation of phages and their regulatory elements in phage therapy and biotechnological applications, respectively.

## 1. INTRODUCTION

As of August 2022, over 600 complete genome sequences of *Pseudomonas* phages, mostly infecting *P. aeruginosa*, are available through the United States National Center for Biotechnology Information (NCBI). These phages display extensive morphological and genomic diversity, and have distinctive replication strategies (Ceyssens & Lavigne, 2010; Ha & Denver, 2018; Sepúlveda-Roble et al., 2012). The majority of these phages belong to the *Caudoviricetes*, the class of tailed, double-stranded DNA phages. Among these, the *Autographiviridae* family currently encompasses nine subfamilies and 133 genera (4). The hallmark feature of the members of this family is that they encode a single-subunit RNA polymerase (RNAP), enabling the phage to self-transcribe its own genes. In addition, all *Autographiviridae* phages display considerable synteny across their genomes.

Escherichia virus T7, one of the best studied bacterial viruses and type species for the *Studiervirinae* subfamily (*Teseptimavirus* genus), was instrumental in unravelling the molecular underpinnings of DNA replication and contributed greatly to the development of molecular biology in general (5). Specifically, its unique transcriptional scheme has become an essential tool in the field of microbial synthetic biology (6). The T7 RNA polymerase and its cognate promoter sequence are used to design highly controlled genetic circuits and drive the expression of desired gene products in various hosts.

T7-like phages have long been considered to replicate by a strictly lytic lifestyle. However, their distinctive genome organisation was found as a prophage in *Pseudomonas putida* KT2440, suggesting that certain *Autographiviridae* phages might be prone to a lysogenic lifestyle (7). Indeed, some T7-like phages have been identified that encoded a site-specific integrase in close proximity of the RNA polymerase gene (8–14). It was only recently demonstrated that certain T7-like phages that infect *Pelagibacter* and *Agrobacterium* do lysogenize their host cells (14, 15).

We here present a novel phage targeting *P. aeruginosa*, LUZ100, a distant relative of the *Teseptimavirus* genus among the *Autographiviridae* family that harbours genes associated with a temperate lifestyle. This phage is, to the best of our knowledge, the first representative of its kind that can infect the pathogen *P. aeruginosa*. We characterised its morphological and biological properties, performed a genomic analysis and charted the LUZ100 transcriptional landscape using the recently established ONT-cappable-seq method (16). This nanopore-based RNA sequencing method enables full-length profiling of primary prokaryotic transcripts and provides a global map of key viral regulatory sequences involved in transcriptional reprogramming of the host cell.

## 2. MATERIALS AND METHODS

### 2.1 Bacterial strains and culture conditions

The collection of *P. aeruginosa* strains in this study includes laboratory strains PAO1k, PA7 and PA14, 24 strains from the Pirnay collection and 47 clinical strains that were isolated from the respiratory tract of cystic fibrosis (CF) patients in the Leuven University Hospital, Leuven, Belgium (17). Bacteria were cultivated in Lysogeny Broth (LB) liquid medium with shaking or plated onto LB agar (1.5% (w/v)) at an incubation temperature of 37°C.

### 2.2 Phage isolation, propagation, and purification

Clinical *P. aeruginosa* strain PaLo41 was used for phage isolation and propagation. Phage LUZ100 was isolated from a sewage sample of the Leuven University Hospital, Leuven, Belgium. The sewage sample was centrifuged at 4,000 *g* for 30 min and filtered using a 0.45 µm filter to remove bacteria and environmental debris. A 5 mL aliquot of filtered water sample was mixed with an equal volume of 2X LB liquid medium, 100 µL of exponentially growing bacterial culture and 1 mM of CaCl_2_, and cultured at 37°C overnight with constant shaking. After adding a few drops of chloroform and centrifugation for 1 hour at 4,000 *g*, the supernatant was filtered again through a 0.45 µm filter and screened for the presence of phages according to the standard double-agar overlay method using a 0.5% LB agar top layer (18). Single plaques were picked up to start a next round of propagation. This process was repeated several times to produce a homogenous phage stock. Next, the phage was amplified on plates by performing five agar overlay assays in parallel and overnight incubation at 37°C. The lysed top layers were collected in a tube, complemented with 30 mL of phage buffer (8.77 g NaCl, 2.47 g MgSO_4,_ 1.21 g Tris-HCl, pH 7.5 in 1 L of dH_2_O) and chloroform (1% v/v) and vigorously shaken overnight. Next, the solution was centrifuged for 40 min at 4,000 *g* and filtered (0.45 µm) to remove residual bacterial debris. The crude phage lysate was concentrated and purified by PEG8000 precipitation as previously described by Ceyssens *et al*. (2006), with minor modifications. Briefly, phages were precipitated overnight at 4°C, spun down at 4,000 *g* for 40 min, resuspended in 3 mL phage buffer and stored at 4°C for further analysis.

### 2.3 Host range analysis

The host range of phage LUZ100 against our characterised collection of *P. aeruginosa* strains was determined by spotting 1.5 µL of 1:10 dilutions of the phage stock on an initiated bacterial lawn of each strain, created with a double agar overlay using a 0.5% LB agar top layer. After overnight incubation at 37°C, the success of phage infection was assessed by surveying the clearance of the spots in the bacterial lawn, and scored as (1) for complete lysis, (2) for lysis with individual plaques, and (3) for no lysis. The bacterial strains were only considered to be susceptible for the phage when distinctive plaques could be observed, as clearing zones could also arise due to lysis effects that do not rely on productive phage infection (20).

### 2.4 Adsorption assay

To assess the adsorption kinetics of LUZ100 to *P. aeruginosa* PaLo41, an adsorption assay was performed using three biological replicates. PaLo41 was grown in LB medium supplemented with 1 mM CaCl_2_ (Sigma Aldrich) and 1 mM MgCl_2_ (Sigma Aldrich) and grown to the early exponential phase (optical density at 600 nm, OD_600_ = 0.3). At this moment, a sample was taken and plated out to determine the bacterial titer (B). Next, the bacteria were infected with LUZ100 at a multiplicity of infection (MOI) of 0.01 and incubated at 37°C. Subsequently, 100 μl samples were taken 1, 5, 10, 15 and 20 minutes (t) post-infection and directly transferred to an Eppendorf tube with an excess of chloroform to kill the bacteria. For each timepoint sample, the phage titre (P) was determined using the double agar overlay method. Finally, the average phage adsorption constant was calculated using the following formula (18):

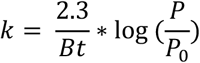

### 2.5 Infection curve

The bacteriolytic activity of phage LUZ100 was determined by monitoring the growth of the phage-infected bacteria over time. For this, overnight cultures of three biological replicates of PaLo41 were inoculated in fresh LB medium and incubated at 37°C to an OD600 of 0.3. Next, these cultures were infected with LUZ100 at an MOI of either 1 or 10. The OD600 of the uninfected and infected cultures was measured every 15 min for 145 minutes on the CLARIOstar® Plus Microplate Reader (BMG Labtech, Ortenberg, Germany) for four technical replicates while incubating at 37°C.

### 2.6 Transmission Electron Microscopy

Transmission Electron Microscopy (TEM) images were made as described by Vallino et al. (2021). Briefly, phage suspension adsorbed on carbon and copper-palladium grids coated with formvar for 3 minutes. Next, the grids were rinsed with water and negatively stained with 0.5% aqueous uranyl acetate. Samples were visualized using a CM10 Transmission Electron Microscope (Philips, Eindhoven, The Netherlands) at a voltage of 80 kV.

### 2.7 LUZ100 receptor analysis

To identify the LUZ100 receptor(s), the genomic DNA (gDNA) of four spontaneous phage-resistant PaLo41 colonies were isolated using the Qiagen DNeasy® Ultraclean® Microbial kit according to manufacturer’s guidelines. the gDNA samples were prepared using the Nextera™ DNA Flex Library Prep kit and paired-end sequencing was performed on an Illumina MiniSeq device. Next, the quality of the reads was assessed with FastQC (v0.11.8) (22) and the adapters and poor-quality reads (phred score <33) were removed from the dataset using Trimmomatic (v0.39) (Bolger et al., 2014). Finally, Snippy (v4.6.0) (Seemann, 2021) was used to identify single nucleotide polymorphisms (SNPs) in the genomes of the LUZ100-resistant PaLo41 clones compared to the PaLo41 reference genome (BioProject PRJNA731114).

### 2.8 LUZ100 genome extraction, sequencing and genomic analysis

LUZ100 phage lysate was subjected to DNase I (Thermo Fisher Scientific) and RNase A treatment (Thermo Fisher Scientific) for one hour in a 37°C water bath. The lysate was treated with sodium dodecyl sulphate (SDS) (Acros Organics), ethylenediamine-tetraacetic acid (EDTA; Sigma Aldrich) and proteinase K (Thermo Fisher Scientific) and incubated for 1 hour in a 56°C water bath. Next, the DNA of LUZ100 was isolated by phenol-chloroform extraction and subsequently purified using ethanol precipitation. The purity and concentration of the DNA was assessed using the SimpliNano™ spectrophotometer (Biochrom US Inc.) and the Qubit™ 4 fluorometer (Thermo Scientific), respectively. Library preparation, Illumina sequencing, and raw read processing was performed as described earlier. The genomic sequence of LUZ100 was assembled with SPAdes (v3.13.2) using default parameters (24), followed by visual inspection of the assembly in Bandage (Wick et al., 2015). In addition, the phage genome was sequenced in full-length using nanopore sequencing to identify the genomic termini. For this, the phage DNA was prepared by the Rapid Barcoding kit (SQK-RBK004) (Oxford Nanopore Technologies) and sequenced on a MinION device (FLO-MIN 106, R9.4). The raw sequencing data was basecalled using Guppy (v3.4.4) and reads were cleaned with Porechop (v0.2.3). The phage genome was assembled using Unicycler (v0.4.8) (26) and subsequently annotated using the RAST pipeline in PATRIC (v3.6.1) (27) by applying the ‘Phage’ annotation recipe (Mcnair et al., 2018; 2019). Manual curation was performed by scanning each predicted CDS for homologs using HMMR, BLASTP and HHPred with default settings, as provided by the MPI Bioinformatics Toolkit (30). This annotation was subsequently used for transcriptomic data analysis. The genome sequence of *Pseudomonas* phage LUZ100 was deposited in NCBI Genbank (BioProject accession number PRJNA870687).

### 2.9 RNA extraction, sequencing and transcriptomic analysis

#### 2.9.1 RNA sampling and extraction

An overnight culture of PaLo41 was diluted 1:100 in 50 ml LB medium, incubated at 37°C and grown to an OD_600_ of 0.3. At this moment, a 4.5 ml sample was mixed with 0.9 ml stop mix solution (95% v/v ethanol, 5% v/v phenol, saturated pH 4.5) and immediately snap-frozen in liquid nitrogen (sample t_0_). The remaining culture was infected with LUZ100 (MOI = 10) and incubated at 37°C. Additional samples were taken every 4 minutes up to 40 minutes after infection. Samples were thawed on ice, pelleted by centrifugation (20 min, 4°C, 4,500 rpm) and resuspended in a 0.5 mg/mL lysozyme solution. Subsequently, all samples, aside from t_0_, were pooled in equal amounts and homogenized (sample t_ϕ_). The resulting samples t_0_ (uninfected) and t_ϕ_ (phage infected) were subjected to hot phenol treatment and ethanol precipitation to extract and purify the RNA. Next, t_0_ and t_ϕ_ were treated with DNase I, and subsequently purified using ethanol precipitation and spin-column purification. The absence of genomic DNA was verified by PCR using a phage-specific and host-specific primer pair (Supplementary Table S1). Finally, the integrity of the RNA samples was evaluated by running the samples on an Agilent 2100 Bioanalyzer using the RNA 6000 Pico Kit. Samples with an RNA integrity number (RIN) ≥ 9 were used for downstream processing and sequencing.

#### 2.9.2 ONT-cappable-seq and data analysis

Prokaryotic RNA samples t_0_ and t_ϕ_ were supplemented with 1 ng of an *in vitro* transcribed control RNA spike-in (1.8 kb), which was synthesized using the HiScribe T7 High Yield RNA Synthesis Kit according to manufacturer’s guidelines (New England Biolabs). Next, library preparation of the RNA samples was performed according to the ONT-cappable-seq method (Putzeys et. al, 2022). Equimolar amounts of the resulting t_0,enriched_, t_0,control_, t_ϕ,enriched_, and t_ϕ,control_ cDNA samples were pooled in a 10 µL volume. The final library was loaded on a MinION flowcell (FLO-MIN 106, R9.4) and sequenced using the MinION platform for >48h until all pores were exhausted. In parallel, the MinIT device with build-in Guppy software (v3.2.10) was used in high-accuracy mode to simultaneously base-call and demultiplex the reads, retaining only the reads with sufficient quality (>7). The overall performance of the sequencing run and raw read quality was assessed using NanoComp (v1.11.2). Next, raw reads were processed and subsequently mapped to the genomes of *Pseudomonas* phage LUZ100 and *P. aeruginosa* host strain PaLo41, as described previously (16). Sequencing quality, read lengths and mapping metrics are reported in Supplementary Table S2. Alignments were visualized in Integrative Genomics Viewer (IGV) (31). Finally, data analysis was performed according to the ONT-cappable-seq workflow (https://github.com/LoGT-KULeuven/ONT-cappable-seq) to identify viral transcriptional start sites (TSS) and termination sites (TTS) and elucidate the transcriptional architecture of LUZ100 (16). To discriminate between phage-specific and host-specific promoter sequences, regions upstream the annotated TSSs (−100 to +1) were analyzed using MEME and the *Pseudomonas* σ70 promoter prediction tool SAPPHIRE.CNN (32, 33). The motif of the identified phage-specific promoter sequences was used to conduct a MAST search on the LUZ100 genome to detect TSS originally missed in the ONT-cappable-seq workflow (34). Based on the MAST results and manual inspection of the LUZ100 transcriptional landscape, three additional phage promoters were added to the list of regulatory elements. In parallel, for each TTS identified with ONT-cappable-seq, the surrounding −60 to +40 region was uploaded in ARNold to predict intrinsic, factor-independent transcription termination sequences (35). The transcription units (TUs) of LUZ100 were defined by neighboring TSS and TTS determined in this study, if supported by ONT-cappable-seq reads that span the candidate TU. In case transcripts from a specific TSS lacked a defined TTS, the longest transcript was used to delineate the TU. Genetic features located on the same strand annotated into a TU in case they were covered by at least 90% (bedtools intersect - F 0.9 -s). Based on the TUs and their gene content, complex operon structures were delineated by finding overlapping TUs with the same orientation that share at least one annotated gene (16, 36).

### 2.10 *In vivo* promoter activity assay

To validate the activity of host-specific promoter P3 *in vivo*, the SEVAtile DNA assembly method was used to clone the promoter upstream of a standardized ribosomal binding site (BCD2) and an *msfGFP* gene (37). The construct was transformed to *E. coli pir2* cells, which were plated on LB agar containing kanamycin (50 µg/ml). A pBGDes vector lacking a promoter (pBGDes BCD2-msfGFP) and a vector containing a constitutive promoter (pBGDes Pem7-BCD2-msfGFP) were used as a negative and positive control, respectively. The used vectors, primers and tiles are listed in Supplementary Table S1. Next, three biological replicates of the transformed *E. coli* cells were inoculated in M9 medium (1x M9 salts (BD Biosciences), 0.5% casein amino acids (LabM, Neogen), 2 mM MgSO4, 0.1 mM CaCl2 (Sigma Aldrich), 0.2% citrate (Sigma Aldrich)) complemented with 50 µg/ml kanamycin. The next day, cultures were diluted in fresh M9 medium, transferred to a Corning® 96 Well Black Polystyrene Micro-plate with Clear Flat Bottom and incubated shaking. After 3.5 hours, the OD_600_ and msfGFP levels were measured on a Clariostar® multimode plate reader (BMG Labtech, Ortenberg, Germany). Then, the msfGFP levels were normalized for OD_600_ and converted to absolute units using 5(6)-carboxyfluorescein (5(6)-FAM) (Sigma Aldrich) as a calibrant (36, 38). Finally, the statistical software JMP® was used to analyse the data (39).

## 3. RESULTS AND DISCUSSION

### 3.1 Biological characteristics of phage LUZ100

#### 3.1.1 Phage morphology and host range

Phage LUZ100 was isolated from hospital sewage water using clinical *P. aeruginosa* strain PaLo41 as host bacterium. After culturing LUZ100 on a bacterial lawn of PaLo41, the phage produces plaques of 4-5 mm in size, surrounded by an opaque-looking halo zone of approximately 1-2 mm (Fig. 1A). Transmission electron microscopy analysis confirms that LUZ100 is a podovirus with a short stubby tail attached to an icosahedral head (Fig. 1B).

**Figure 1.**
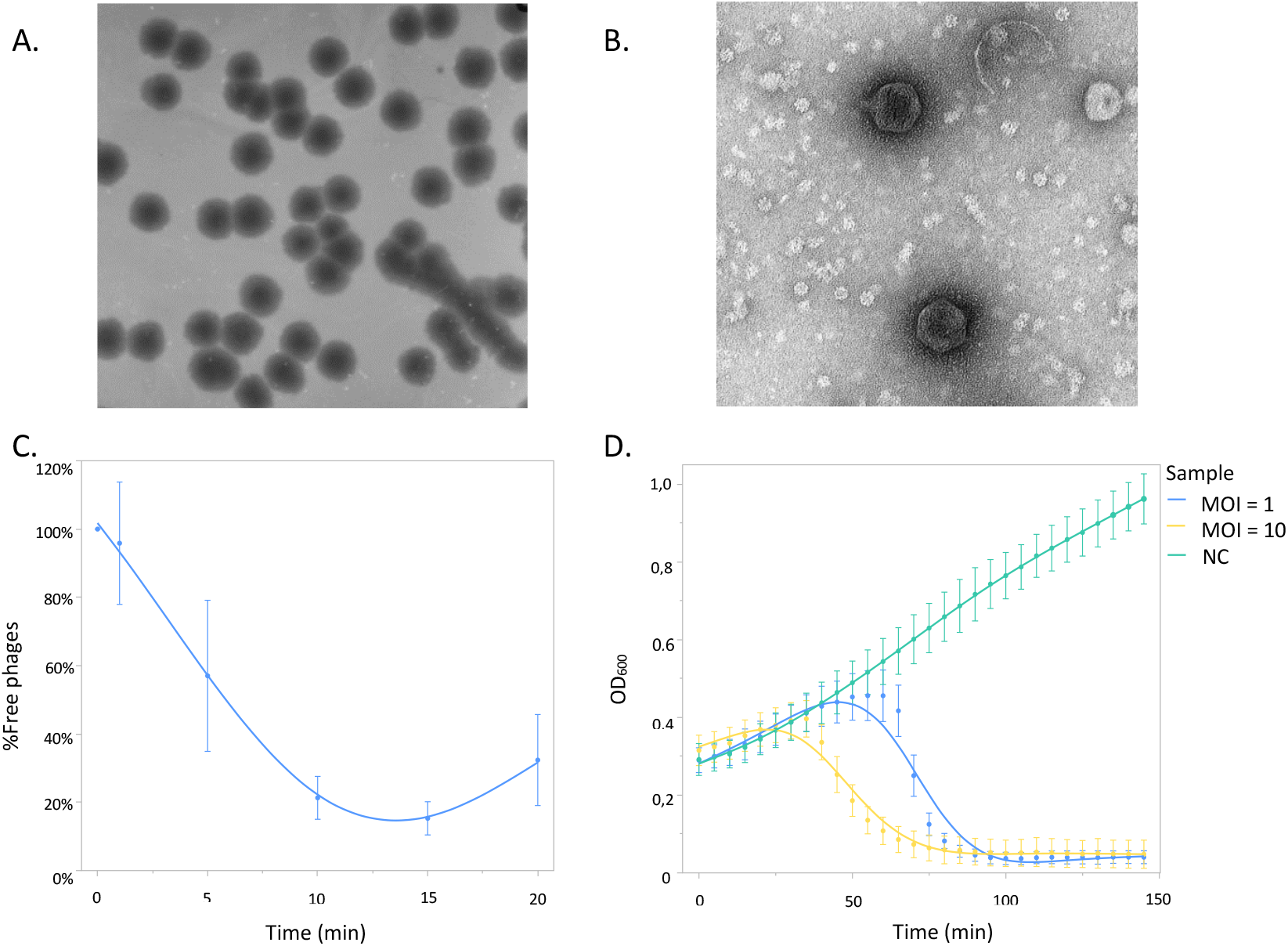
Morphological and microbiological features of phage LUZ100. (A) Plaques of LUZ100 on *P. aeruginosa* PaLo41, showing large bulls-eyed plaques (B) Transmission electron microscopy image demonstrating the podovirus virion morphology of LUZ100. (C) Adsorption curve of LUZ100 on *P. aeruginosa* PaLo41, showing that 85% of the phages is adsorbed to the host cell surface after 15 minutes. (D) Infection curves of LUZ100 infecting *P. aeruginosa* PaLo41. At MOI 10, LUZ100 completes its infection cycle after approximately 30 minutes causing lysis of the bacterial cells. (MOI = multiplicity of infection, NC = Negative Control, OD_600_ = optical density at 600 nm).

The host range and lytic activity of the phage was determined on a diverse collection of 74 *P. aeruginosa* isolates, including 71 clinical strains and the laboratory strains PAO1k, PA7 and PA14. Phage LUZ100 displays a narrow host spectrum, infecting 24% of the strains in the panel (Supplementary Fig. S1).

#### 3.1.2 Phage adsorption rate

An adsorption assay was performed to assess the efficiency and timing of LUZ100 phage infection. After 15 minutes, 85% of the phages were adsorbed to the host cell surface (Fig.1C). The average adsorption constants (k) after one and five minutes equal 8.00*10^−9^ ml/min and 2.63*10^−9^ ml/min, respectively. In comparison to other members of the T7-like phages, such as *P. putida* phage phi15 (k = 2.51*10^−8^ ml/min after one minute) and *P. fluorescens* phage IBB-PF7A (k = 5.58*10^−10^ ml/min after five minutes), LUZ100 has a moderate adsorption rate (40, 41).

#### 3.1.3 Infection curves

LUZ100 infection curves of PaLo41 infected at different MOIs were analysed to determine when bacteria begin to lyse after phage infection (Fig. 1D). At MOI 10, the optical density decreases after approximately 30 minutes, marking the completion of the LUZ100 infection cycle. In addition, the virulence of LUZ100 was assessed by calculating the virulence index and phage score. These metrics equal 0.069 and 0.065 respectively, which is low in contrast to the strictly lytic phage T7 (virulence index of 0.84) (42, 43). The reduced LUZ100 lytic activity is logically related to its potential temperate character, discussed below. However, attempts to generate stable lysogens have not been successful so far (data not shown).

#### 3.1.4 Identification of phage receptor

Whole-genome sequencing of four spontaneous phage-resistant PaLo41 mutants was carried out to identify the bacterial receptor to which LUZ100 binds to its host. When compared to the host genome, two out of the four mutants showed SNPs in the coding sequence of O-antigen ligase (WaaL) (Supplementary Table S3). WaaL is involved in the Lipopolysaccharide (LPS) biosynthesis pathway (Abeyrathne et al., 2005), and hence, we propose that LPS is a likely receptor for phage LUZ100. Consistent with these results, LPS is recognized by several other members of the T7-like phages, including gh-1 and T7 but differs from other known T7-like *Pseudomonas* phages such as phiKMV and LUZ19 that use type IV pili as primary receptors (44–47).

### 3.2 Genomic and transcriptomic analysis of LUZ100

#### 3.2.1 General genomic features of LUZ100

LUZ100 has a linear double-stranded DNA genome containing 37,343 base pairs (bp), flanked by 244 bp direct terminal repeats. The genome has an average GC-content of 61% and encodes 56 open reading frames (ORFs) and two tRNAs (Supplementary Table S4). The biological function of 30 translated ORFs could be predicted, leaving 26 ORFs assigned as hypothetical proteins. The predicted features and overall genomic organisation of phage LUZ100 reveal remarkable similarity to members of the *Studiervirinae* in the *Autographiviridae* phage family, including coliphage T7 (NC_001604) and *Pseudomonas* phage gh-1 (NC_004665.1) (44) (Fig. 2A). In analogy with the canonical T7 gene order, the LUZ100 genome can be roughly organized into three functional and temporal modules, representing the genomic regions involved in host takeover (early), DNA metabolism (middle) and virion morphogenesis (late) (48). However, at the nucleotide level, the genomic sequence of LUZ100 is unique, showing only distant sequence similarity to other phages when queried against the NCBI *Caudoviricetes* database (blastn query coverage <10%; Per. Identity <80%). In addition, unlike other T7-like phages infecting *P. aeruginosa*, the LUZ100 genome encodes an integrase gene (tyrosine site-specific recombinase) upstream of the RNA polymerase gene, suggesting that this phage could lysogenize its host cells (14, 15).

**Figure 2.**
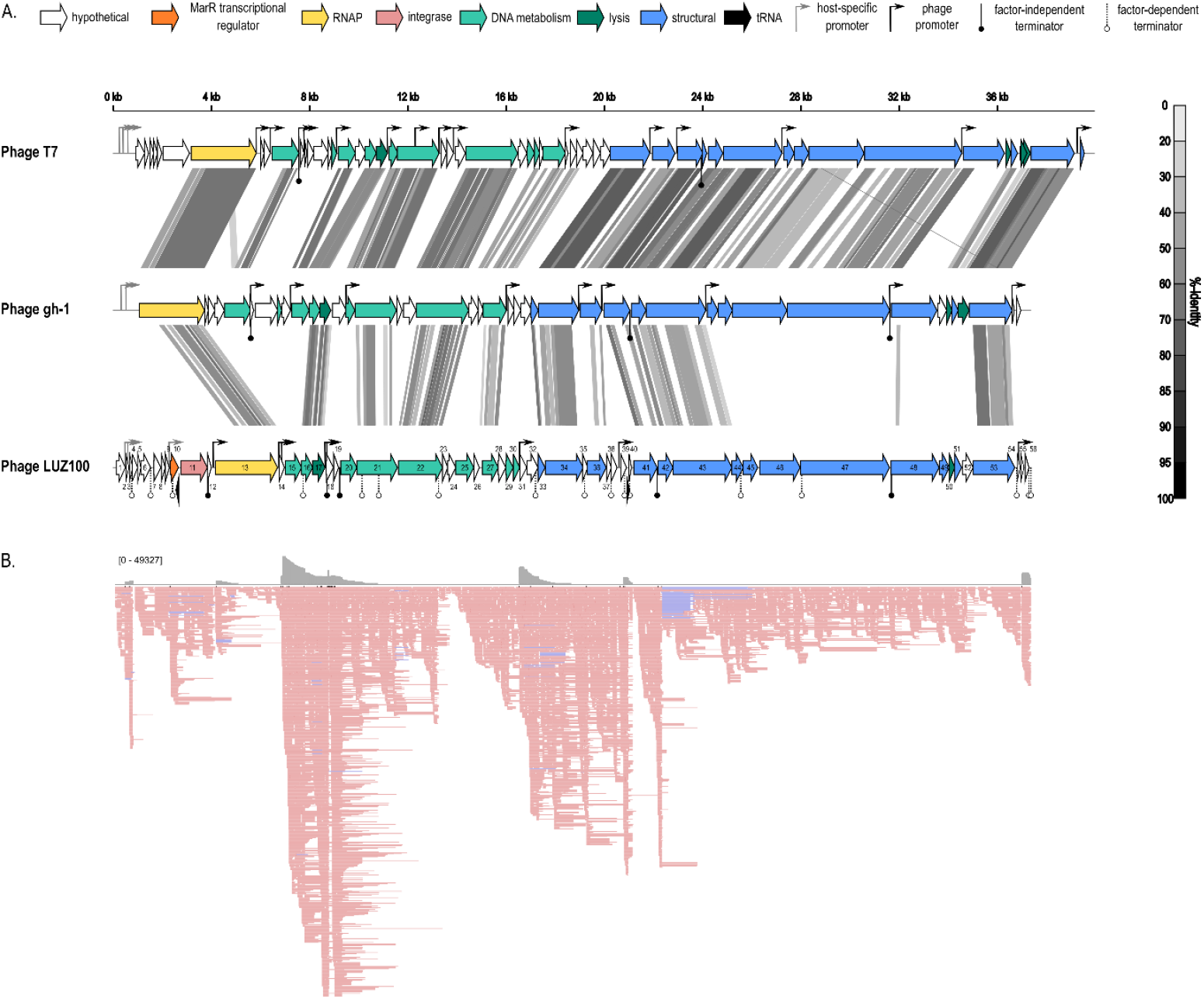
Genomic and transcriptomic overview of *Pseudomonas* phage LUZ100. (A.) comparison of the genomes of phage T7, gh-1 and novel *Pseudomonas* phage LUZ100 are schematically depicted, each arrow representing an ORF. The tRNA genes (black), phage RNAP (yellow), and genes involved in DNA metabolism (green), virion structure (blue), lysis (dark blue) and integration (pink), are highlighted in different colours. Pairwise genome comparisons were generated by tBLASTx in VipTree (49) and show the %-identity of similar regions in greyscale. Promoters are indicated with rightward arrows, where the host-specific promoters (grey) and phage RNAP-specific promoters (black) are marked in different colours. Circles below the genomic map represent terminators, with putative factor-independent terminator sequences indicated in black. (B.) ONT-cappable-seq data track of the transcriptomic landscape of phage LUZ100 visualized in IGV (downsampling with 10 bases window size, 50 of reads per window). The upper part represents the coverage plot and the lower part visualizes the read alignments. Reads that align to the Watson strand and Crick strand are indicated in pink and blue, respectively.

#### 3.2.2 The transcriptional landscape of LUZ100

To gain more insights in the transcriptional scheme and gene regulation mechanisms of T7-like phages with temperate elements, we studied the viral transcriptome of LUZ100 using ONT-cappable-seq. In contrast to classic RNA sequencing approaches, ONT-cappable-seq allows end-to-end sequencing of primary transcripts and the identification of key regulatory elements in dense phage transcriptomes, such as transcription initiation and termination sites, as well as operon structures (16). To explore the phage transcriptome in a global fashion, RNA from multiple timepoints throughout LUZ100 infection of its *P. aeruginosa* PaLo41 host were pooled together and sequenced on a MinION sequencing device.

ONT-cappable-seq data analysis, followed by manual curation, revealed a total of 11 viral transcription start sites (TSS) and 22 transcription termination sites (TTS), together with their associated promoter (Table 1) and terminator sequences (Supplementary Table S5), across the LUZ100 genome (Fig. 2). Among the transcription termination sequences, five are predicted to be intrinsic, factor-independent terminators. Notably, many of the regulatory elements in LUZ100 are in conserved positions relative to T7, suggesting that the transcriptional scheme of LUZ100 is coordinated in a similar manner to the type coliphage.

**Table 1.**
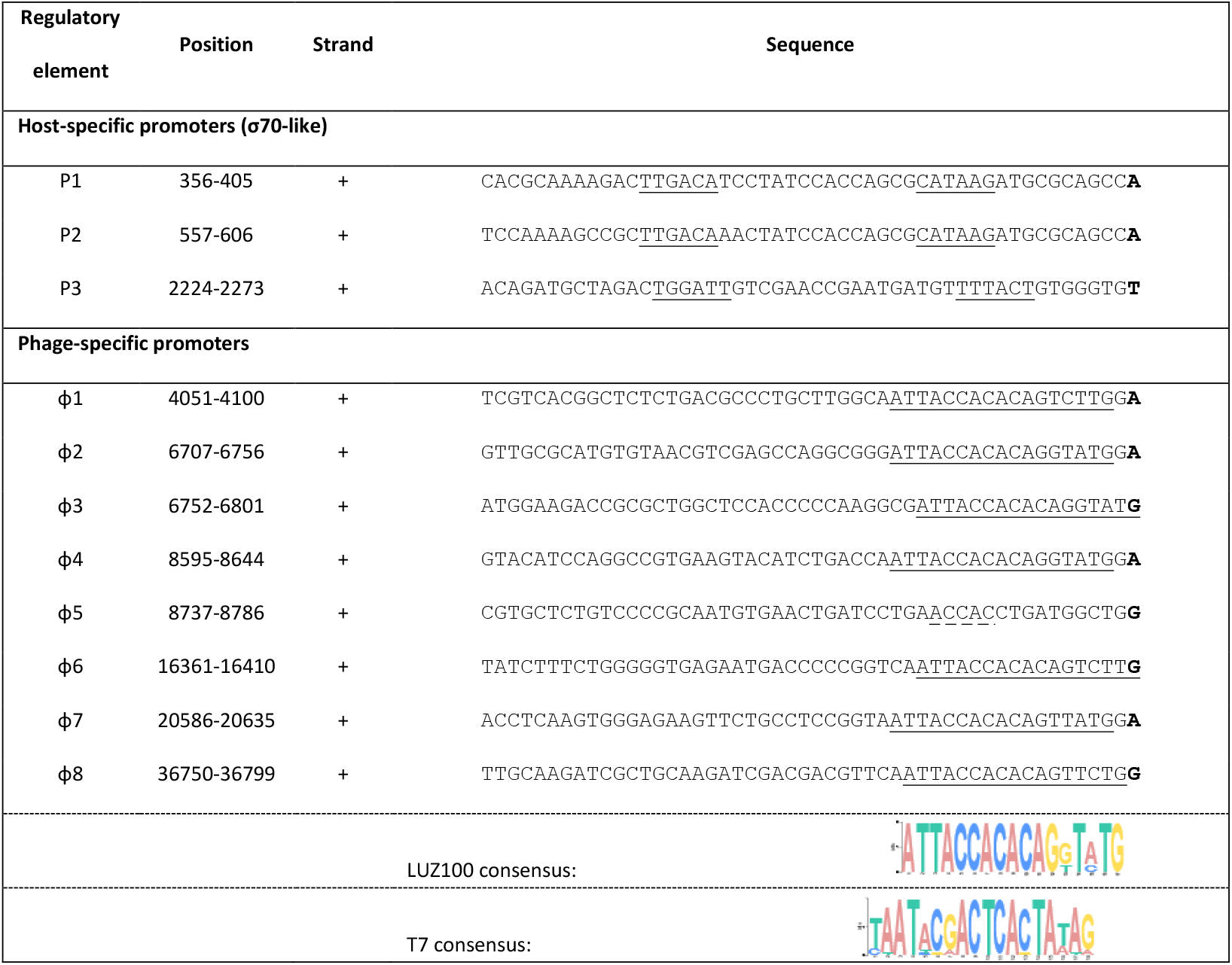
Overview of LUZ100 promoter elements identified by ONT-cappable-seq. TSSs are indicated in bold. In case of the host-specific phage promoters, the σ70-like sequences are underlined. For the phage-specific promoters, a consensus motif was found using MEME (v 5.4.1) and is marked with the black line.

In agreement with most *Autographiviridae* phages, the early genomic region of LUZ100 encodes a suite of hypothetical proteins that are presumably involved in the host takeover process. In this region, we identified three TSSs and their cognate promoter sequences that show significant similarity to the σ70 consensus sequence of *Pseudomonas*. Upon infection, these host-specific promoters (P1, P2, and P3) are likely to be recognized by the bacterial RNAP and drive the expression of early genes. Interestingly, promoter P3, which drives transcription of the putative lysogeny gene module of LUZ100, deviates from the strongly conserved σ70 sequences of promoter P1 and P2. In addition, we observe that the majority of the transcripts that start at P3 are strongly terminated by terminator T4, located directly downstream of the integrase gene. These findings suggest that transcription of the lysogeny-associated genes is regulated in a distinct, independent manner from the other early genes, which is explored in more depth below.

The end of the early genomic region encodes the DNA-directed RNA polymerase. This RNAP shows high sequence similarity to the well-studied T7 RNAP (blastp e-value of 9e^−173^). However, the LUZ100 RNAP domains that specifically recognise and bind the phage promoter sequences, are only remotely related to the corresponding RNAP regions of phage T7 and various T7-like *Pseudomonas* phages (Supplementary Table S6) (50, 51). This hints towards an altered promoter specificity between the LUZ100 RNAP and the RNAP of T7 group members that infect *Pseudomonas* hosts. Based on the ONT-cappable-seq data, we identified five additional promoter sequences that are likely to be specifically recognized by the LUZ100 RNAP. These promoters share a highly conserved 17 bp motif (e-value = 4.7*10^−10^) that only partially resembles the T7 consensus promoter (5’-TAATACGACTCACTATA**G**) (Table 1) (52). This motif was used to manually recover three additional promoters from the transcriptomic data that were not included previously, as they did not meet the stringent TSS threshold value. This yielded a total of eight phage-specific promoter sequences with a distinctive motif, further supporting the hypothesized orthogonality between different T7-like phage RNAP-promoter pairs (53).

The middle and late genes, which are transcribed by the phage RNA polymerase from their cognate phage promoters, are mainly responsible for phage DNA metabolism and virion morphogenesis, respectively (48). LUZ100 is equipped with the hallmark T7-like replication machinery, including a predicted single-stranded DNA binding protein, an endonuclease, a DNA polymerase, a DNA primase/helicase, and an exonuclease. In addition, the phage encodes a lysozyme-like protein that resembles the well-studied T7 lysozyme, which is involved in host cell lysis and inhibition of T7 RNAP transcription activity (blastp expected value of 3e^−36^) (Cheng et al., 1994; Huang et al., 1999). In LUZ100, expression of genes involved in DNA metabolism appears to be largely driven by phage promoters ϕ2, ϕ3, ϕ4 and ϕ5. Remarkably, promoters ϕ2 and ϕ3, are organized in tandem, directly upstream of an annotated single-stranded DNA-binding (SSB) protein. A similar observation was made for N4-like *Pseudomonas* phage LUZ7, in which two promoters in tandem achieved extremely high expression levels of a key SSB protein, the transcriptional regulator Drc (56). These results suggest that tandem promoters might be a common theme in phages to drive the expression of SSB proteins, which are often required in high abundance.

In contrast to T7, the replication module of LUZ100 contains three additional genes that show homology to a thymidylate synthase, a ribonucleoside-diphosphate reductase (RNR), and a nucleotide pyrophosphohydrolase-like protein of the MazG protein family. These genes were presumably acquired from the host by horizontal gene transfer (15). Interestingly, auxiliary metabolic genes (AMGs), including RNRs and thymidylate synthases, were also found in other T7-like temperate phage genomes and are generally thought to reinforce the host metabolism during the phage infection process (13, 15). Both RNRs and thymidylate synthases are known to be involved in DNA metabolism and can presumably facilitate phage replication in the infected cell by increasing deoxynucleotide biosynthesis (57, 58). In contrast, phage-encoded nucleotide pyrophosphohydrolases are hypothesized to play a role in the regulation of cellular (p)ppGpp levels to maximise infection efficiency (59, 60).

Finally, the late genomic region of LUZ100 mainly encodes proteins involved in virion structure, assembly, DNA packaging and host cell lysis. The structural gene cassette of LUZ100 includes genes coding for the portal protein, the major capsid protein (MCP), the tail fibre protein, tail tubular proteins A and B, internal virion proteins and the small and large terminase subunits. In analogy with T7, gene expression of the LUZ100 MCP appears to be tightly controlled by a local phage-specific promoter and an apparent strong terminator sequence located immediately downstream of the MCP gene. It should be noted that no obvious T7 internal virion proteins C (gp15) and D (gp16) equivalents could be detected based on amino acid sequence similarity. However, given the strict synteny of the structural modules of T7, gh-1 and LUZ100, the translated products of LUZ100 *gp46* and *gp47* are likely to be functionally related. The ONT-cappable-seq data also revealed interesting transcriptional activity downstream of the LUZ100 MCP. Part of the transcripts in the structural region of the phage were transcribed antisense. A tBLASTx search was performed to verify whether incomplete annotation could explain the presence of this unexpected transcriptional activity. However, no obvious protein coding sequences were identified, leading us to speculate that the antisense transcripts correspond to a non-coding antisense RNA molecule. As previously hypothesized for LIT1 and LUZ7, these antisense RNAs putatively have a regulatory role in expression of the structural proteins (16, 61).

At the end of the infection cycle, the newly synthesized phage progeny will be released by lysing the host cell. In general, the lysis pathway of T7-like phages is largely mediated by three elements: lysozyme, holin (type II) and spanins (Rz/Rz1), all targeting different layers of the bacterial cell envelope. The LUZ100 lysozyme and holin could be identified and are actively being transcribed during infection. However, no apparent spanin gene equivalents could be annotated after screening the phage ORFs for membrane localisation signals that hallmark the internally overlapping Rz/Rz1 gene pair, as observed in T7 relatives (62). It has been suggested that, under certain physiological conditions, the lack of spanins could impede phage-mediated lysis and subsequent progeny release from Gram-negative bacteria (63). However, under laboratory conditions, LUZ100 appears to successfully breach the cell barriers of its host.

#### 3.2.3 LUZ100 transcription unit architecture

Besides pinpointing transcriptional landmarks, the ONT-cappable-seq data was also used to elucidate the transcription unit architecture of LUZ100. Based on adjacent pairs of TSS and TTS defined in this work, we identified 37 unique transcription units (TUs) that cover on average 3.6 genes (Supplementary Table S7). This large number of TUs is likely to be the result of sequential read-through across different TTSs (16). In general, we found that the genes that are co-transcribed in a TU are functionally related, as expected. For instance, whereas TU4 is devoted to transcription of the lysogeny module, TU17 and TU31 span genes involved in DNA metabolism, and virion morphology, respectively. In addition, several TUs show significant overlap in terms of their gene content, suggesting that overlapping TUs provide an alternative mechanism to finetune the expression levels for specific genes. Similar observations have been made in bacteria, where complex TU clusters are thought to be employed as a regulatory strategy to modulate gene expression levels under different conditions (64–66).

### 3.3 Hypothesized lysogeny of LUZ100

Recently, multiple incidences of temperate T7-like phages have been reported, contradicting the general assumption that T7-like phages propagate according to a strictly lytic life cycle (14, 15, 67, 68). However, general phage characterisation of LUZ100, in combination with genomic and transcriptomic analyses, revealed low virulence and an actively transcribed phage-encoded integrase. These traits are both clues that LUZ100 is prone to a lysogenic life cycle and put LUZ100 forward as a new member of the emerging clade of T7-like temperate *Autographiviridae* phages.

Another feature that points in the same direction is the presence of two tRNA genes encoded on the LUZ100 genome. tRNAs are abundantly present in phage genomes. However, when virulent and temperate phages are compared, the former tend to have more copies (69). In addition to the low copy number of tRNAs encoded in LUZ100’s genome, one of the tRNA genes, a tRNA-Leu, is located immediately upstream of the integrase gene. Since tRNAs are considered integrational hotspots for phages and other mobile genetic elements, the origin of this tRNA molecule might be recruited for imprecise excision after a previous integration event (70). A similar hypothesis was proposed for temperate T7-like *Pelagibacter* phages that also retained a tRNA-Leu sequence upstream of their integrase (15). The remnants of the previous integration event in combination with the low tRNA abundance in LUZ100’s genome are additional clues that link LUZ100 to a lysogenic lifecycle.

As previously mentioned, transcriptome analysis revealed that the LUZ100 integrase gene is actively transcribed. Directly upstream of the integrase, LUZ100 encodes a MarR-like transcriptional regulator. This regulator has been identified in several integrase-coding T7-like phages and was recently proposed to be involved in their lytic/lysogeny decision (8, 14, 15). The ONT-cappable-seq data shows that the MarR-like regulator and the integrase are co-transcribed in an operon from host-specific promoter P3. Since this promoter deviates from the highly conserved σ70 sequences of promoter P1 and P2, a fluorescence expression assay was performed to validate the activity of the promoter *in vivo*. For this purpose, the P3 promoter was cloned upstream of a ribosomal binding site (RBS) and an *msfGFP* gene using the SEVAtile DNA assembly method. Subsequently, the resulting vector was transformed to the laboratory model organism *E. coli* (37). The P3 promoter shows significantly (p-value < 0.001) higher expression of the fluorescent reporter compared to the negative control without promoter sequence, confirming its activity *in vivo* (Supplementary Fig. 2). The transcriptional activity of both the MarR-like regulator and the integrase further supports our hypothesis that LUZ100 is prone to a lysogenic lifecycle through MarR-based regulation.

The phage RNAP is key to complete a lytic infection cycle, and the regulation of the polymerase is proposed to be intertwined with the MarR-based transcriptional regulation in some of the temperate T7-like phages (14). During lysogeny, the MarR regulator is hypothesized to repress phage RNAP expression through recognition of specific binding sites sequences that flank the promoter sequence upstream the RNAP (14). Interestingly, in case of LUZ100, the location of phage-specific promoter ϕ1, encoded directly upstream of the RNAP, raises questions on how the RNAP becomes initially activated. Visual inspection of the long-read transcriptional landscape of the lysogeny cassette and RNAP of LUZ100, revealed that not all transcripts are terminated after the integrase gene. Based on these findings, we hypothesize a lysogeny control mechanism where the host RNAP sporadically reads through terminator T4 and transcribes the phage RNAP from host-specific promoter P3, located further upstream (Fig. 3A). Once the phage RNAP is transcribed by the host RNAP and expressed, it will be able to bind the phage-specific promoter and induce its own transcription in a positive feedback loop (Fig. 3B). In contrast, when the lysogenic state is favoured, transcription from the phage-specific promoter is repressed by the MarR-like protein, and middle and late gene expression is impaired (Fig. 3C). However, it should be noted that our dataset did not capture any sufficiently long reads starting at promoter P3 that span the RNAP entirely, which might be attributed to the limited sequencing depth and technical limitations of ONT-cappable-seq (16).

**Figure 3.**
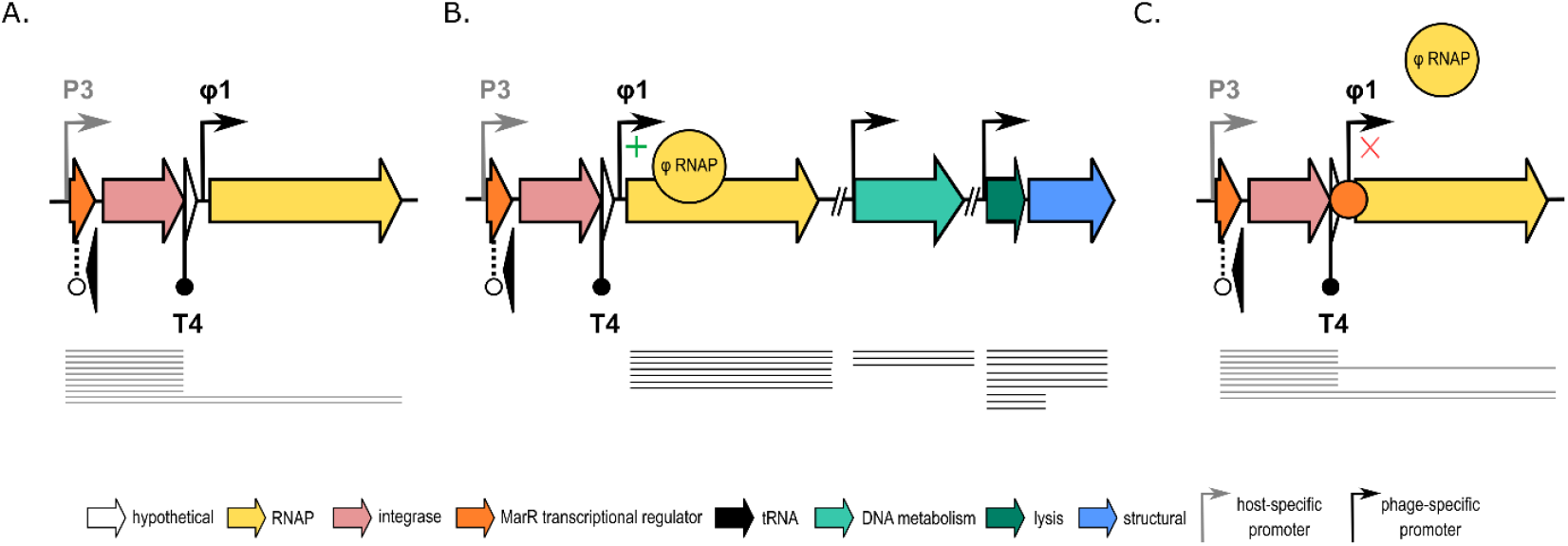
Schematic representation of the hypothesized MarR-based lysogeny control mechanism of LUZ100. (A.) Schematic of putative MarR-based lysogeny control region of LUZ100. Upon infection, the host RNAP transcribes the lysogeny module, including the MarR-like protein and integrase. A limited number of these transcripts read through T4 and transcribe the phage RNAP. (B.) Once expressed, the phage RNAP can initiate transcription from phage-specific promoter ϕ1 and amplify its own transcription. Sufficient expression of the phage RNAP enables expression of the LUZ100 middle and late genes from phage-specific promoters, which are required to complete the lytic infection cycle. (C.) Lysogeny is established by binding of the MarR-like repressor protein to specific binding sites that inhibit transcription initiation from ϕ1 by the phage RNAP.

## 4. CONCLUSION AND PERSPECTIVES

The enormous diversity of bacteriophages is yet again highlighted in our analysis of a novel phage infecting *P. aeruginosa*, LUZ100. In contrast to the strictly lytic model-phage T7, a distant relative among the *Autographiviridae* family, LUZ100 encodes key genes associated with temperate behaviour. Recently, a number of T7-like phages have been identified that show similar characteristics to LUZ100 (14, 15, 67). However, concrete evidence of entering and maintaining the lysogenic lifecycle has remained ambiguous for these phages (14, 15, 67). Also in our research, no stable lysogens could be obtained to date.

To this end, transcriptomics-driven characterisation of bacteriophage LUZ100 was performed to study the molecular processes at work in this peculiar member of the *Autographiviridae* (16). The ONT-cappable-seq data showed active co-transcription of the MarR-like transcriptional regulator and the integrase from the host-specific promoter P3, which further strengthened the hypothesis that not all T7-like phages should straightforwardly be considered as having a strictly lytic life cycle. Moreover, the identification of a phage-specific promoter driving transcription of the phage-encoded RNA polymerase suggests that the regulation of this polymerase is interwoven with the activity of the MarR-based regulator and plays a role in the lytic/ lysogeny switch of LUZ100.

In addition to our findings, others have speculated that from an evolutionary perspective, T7-like phages could be descendants from a common temperate ancestor that (partly) lost its ability to lysogenize its host over time (14). This has several consequences for fields like phage therapy, where virulent phages are preferred for therapeutic purposes (71). Consistently screening for characteristics associated with a lysogenic lifecycle is required to put forward phages suitable for phage therapy. As shown in this work, the combination of multiple omics techniques can serve this purpose. Where DNA-sequencing can identify the presence of lysogeny-related genes, RNA sequencing approaches can give information on their transcriptional activity and underlying regulatory mechanisms.

In addition, the ONT-cappable-seq data allowed us to identify key transcriptional elements of LUZ100, including TSS, TTS and transcription units. The RNAP-promoter pairs identified in this work are distinct from other T7-like phages and could be further explored for synthetic biology applications. Furthermore, other peculiar transcriptional features, such as presence of asRNA and a phage-specific promoter in front of the phage RNAP show that our knowledge on the molecular diversity of phages overall remains limited. However, the ONT-cappable-seq approach has the potential to help bridge these knowledge gaps by generating a bird’s-eye view of phage transcriptomes in an efficient yet cost-effective manner.

## 5. ACKNOWLEDGEMENTS

The authors thank Dr. D. Holtappels for its advice during the experimental work, S. Van Overfelt for helping out with host-range experiments and Dr. J-P Pirnay for providing the collection of clinical *P. aeruginosa* strains. This work was supported by the European Research Council (ERC) under the European Union’s Horizon 2020 research and innovation programme [819800]. JP and MB are funded by a grant from the Special Research Fund [iBOF/21/092].

## 6. DATA AVAILABILITY

The genomes of *P. aeruginosa* strain PaLo41 and phage LUZ100 were deposited in NCBI GenBank (BioProject accession numbers PRJNA731114 and PRJNA870687, respectively). Raw RNA sequencing files are deposited under GEO accession number GSE211961). All scripts and codes used in this study are made available on Github (https://github.com/LoGT-KULeuven/ONT-cappable-seq). Any additional information is accessible from the authors upon request.

Note: private reviewer links to the BioProjects

- PaLo41:

https://dataview.ncbi.nlm.nih.gov/object/PRJNA731114?reviewer=s7u5dra6n7vdbh89o0ia5ai9jh

- LUZ100:

https://dataview.ncbi.nlm.nih.gov/object/PRJNA870687?reviewer=9qd1n4k29i3hsr87amjf6tbi7v

## Supplementary materials

**Supplementary Table S1**.| Overview of the primers, inserts and vectors used in this study.

**Supplementary Table S2**.|Overview of the sequencing metric. (A) Raw reads. (B) Processed reads. (C) Mapped reads.

**Supplementary Table S3**.|Overview of the single nucleotide polymorphisms found in each of the four phage resistant PaLo41 mutants.

**Supplementary Table S4**.| Overview of the LUZ100 coding sequences. Sequence similarity to T7 homologues are indicated in green, while similarity to other blasp hits is represented in black.

**Supplementary Table S5**. | Overview of LUZ100 terminators identified by ONT-cappable-seq. TTS are indicated in bold. Intrinsic, factor independent terminator sequences predicted by ARNold are indicated in blue. Their characteristic stem-loop structure is marked by the blue DNA sequence (stem) surrounding the underlined bases (loop).

**Supplementary Table S6**. | Alignment of the amino acid sequences of T7-like phage RNA polymerase (RNAPs) involved in recognition and binding to promoter sequences.

**Supplementary Table S7**. | Overview of the transcriptional units of LUZ100. The transcriptional units are delineated by the TSS and TTS identified by ONT-cappable-seq.

**Supplementary Figure S1**. | Host range analysis of LUZ100. The susceptibility to LUZ100 phage infection was tested for 47 clinical isolates (PaLo1 – 47), 24 isolates from the Pirnay collection and three reference strains (PAO1k, PA7, and PA14). The light green colour indicates that the phage lysate induced a bactericidal effect on the bacterial lawn, without generating plaques, while the dark green colour shows clearing of the bacterial lawn with the formation of plaques. The pink colour corresponds to a lack of infection.

**Supplementary Figure S2**. | *In vivo* experimental validation of promoter P3 in *E. coli*. The promoter activity was determined by measuring the levels of msfGFP. They were normalized for OD600 and converted into absolute values using 5(6)-carboxyfluorescein (5(6)-FAM) as a calibrant (represented by the 5(6)-FAM/OD600 axis). The negative control (NC) represents a pBGDes vector lacking a promoter (pBGDes BCD2-msfGFP), while the positive control (PC) shows the results for a vector containing a constitutive promoter (pBGDes Pem7-BCD2-msfGFP). The asterisk (*) indicates a significant difference of the promoter in comparison to the NC (based on a Dunnett test, p-value < 0.001). Data represent the mean value of three biological replicates and standard deviation is indicated with error bars.

## REFERENCES

1. Ha AD, Denver DR. 2018. Comparative genomic analysis of 130 bacteriophages infecting bacteria in the genus Pseudomonas. Front Immunol 9:1–13.

2. Ceyssens P-J, Lavigne R. 2010. Bacteriophages of Pseudomonas. Future Microbiol 5:1041–1055.

3. Sepúlveda-Robles O, Kameyama L, Guarneros G. 2012. High diversity and novel species of Pseudomonas aeruginosa bacteriophages. Appl Environ Microbiol 78:4510–4515.

4. Adriaenssens EM, Sullivan MB, Knezevic P, van Zyl LJ, Sarkar BL, Dutilh BE, Alfenas-Zerbini P, Łobocka M, Tong Y, Brister JR, Moreno Switt AI, Klumpp J, Aziz RK, Barylski J, Uchiyama J, Edwards RA, Kropinski AM, Petty NK, Clokie MRJ, Kushkina AI, Morozova V V., Duffy S, Gillis A, Rumnieks J, Kurtböke İ, Chanishvili N, Goodridge L, Wittmann J, Lavigne R, Jang H Bin, Prangishvili D, Enault F, Turner D, Poranen MM, Oksanen HM, Krupovic M. 2020. Taxonomy of prokaryotic viruses: 2018-2019 update from the ICTV Bacterial and Archaeal Viruses Subcommittee. Arch Virol 165:1253–1260.

5. Kutter E, Guttman B. 2001. T Phages, p. 1921–1930. In Brenner, S, H. Miller, J (eds.), Encyclopedia of Genetics, Academic Press.

6. Wang W, Li Y, Wang Y, Shi C, Li C, Li Q, Linhardt RJ. 2018. Bacteriophage T7 transcription system: an enabling tool in synthetic biology. Biotechnol Adv 36:2129–2137.

7. Nelson KE, Weinel C, Paulsen IT, Dodson RJ, Hilbert H, Martins Dos Santos VAP, Fouts DE, Gill SR, Pop M, Holmes M, Brinkac L, Beanan M, DeBoy RT, Daugherty S, Kolonay J, Madupu R, Nelson W, White O, Peterson J, Khouri H, Hance I, Lee PC, Holtzapple E, Scanlan D, Tran K, Moazzez A, Utterback T, Rizzo M, Lee K, Kosack D, Moestl D, Wedler H, Lauber J, Stjepandic D, Hoheisel J, Straetz M, Heim S, Kiewitz C, Eisen JA, Timmis KN, Düsterhöft A, Tümmler B, Fraser CM. 2002. Complete genome sequence and comparative analysis of the metabolically versatile Pseudomonas putida KT2440. Environ Microbiol 4:799–808.

8. Huang S, Zhang S, Jiao N, Chen F. 2015. Comparative genomic and phylogenomic analyses reveal a conserved core genome shared by estuarine and oceanic cyanopodoviruses. PLoS One 10:1–17.

9. Labrie SJ, Frois-Moniz K, Osburne MS, Kelly L, Roggensack SE, Sullivan MB, Gearin G, Zeng Q, Fitzgerald M, Henn MR, Chisholm SW. 2013. Genomes of marine cyanopodoviruses reveal multiple origins of diversity. Environ Microbiol 15:1356–1376.

10. Zhao Y, Temperton B, Thrash JC, Schwalbach MS, Vergin KL, Landry ZC, Ellisman M, Deerinck T, Sullivan MB, Giovannoni SJ. 2013. Abundant SAR11 viruses in the ocean. Nature 494:357–360.

11. Santamaría RI, Bustos P, Sepúlveda-Robles O, Lozano L, Rodríguez C, Fernández JL, Juárez S, Kameyama L, Guarneros G, Dávila G, González V. 2014. Narrow-Host-Range Bacteriophages That Infect Rhizobium etli Associate with Distinct Genomic Types. Appl Environ Microbiol 80:446–454.

12. Pope WH, Weigele PR, Chang J, Pedulla ML, Michael E, Houtz JM, Jiang W, Chiu W, Hatfull GF, Roger W, King J. 2017. Genome Sequence, Structural Proteins, and Capsid Organization of the Cyanophage Syn5: A “Horned” Bacteriophage of Marine Synechococcus 32:736–740.

13. Sullivan MB, Coleman ML, Weigele P, Rohwer F, Chisholm SW. 2005. Three Prochlorococcus cyanophage genomes: Signature features and ecological interpretations. PLoS Biol 3:e144.

14. Boeckman J, Korn A, Yao G, Ravindran A, Gonzalez C, Gill J. 2022. Sheep in wolves’ clothing: Temperate T7-like bacteriophages and the origins of the Autographiviridae. Virology 568:86–100.

15. Zhao Y, Qin F, Zhang R, Giovannoni SJ, Zhang Z, Sun J, Du S, Rensing C. 2019. Pelagiphages in the Podoviridae family integrate into host genomes. Environ Microbiol 21:1989–2001.

16. Putzeys L, Boon M, Lammens E-M, Kuznedelov K, Severinov K, Lavigne R. 2022. Development of ONT-cappable-seq to unravel the transcriptional landscape of Pseudomonas phages. Comput Struct Biotechnol J 20:2624–2638.

17. Pirnay JP, Bilocq F, Pot B, Cornelis P, Zizi M, Van Eldere J, Deschaght P, Vaneechoutte M, Jennes S, Pitt T, De Vos D. 2009. Pseudomonas aeruginosa Population Structure Revisited. PLoS One 4:e7740.

18. Adams MH. 1959. Bacteriophages. Interscience Publishers, Inc., New York.

19. Ceyssens P-J, Lavigne R, Mattheus W, Chibeu A, Hertveldt K, Mast J, Robben J, Volckaert G. 2006. Genomic Analysis of Pseudomonas aeruginosa Phages LKD16 and LKA1 : Establishment of the KMV Subgroup within the T7 Supergroup. J Bacteriol 188:6924–6931.

20. Mirzaei MK, Nilsson AS. 2015. Isolation of Phages for Phage Therapy : A Comparison of Spot Tests and Efficiency of Plating Analyses for Determination of Host Range and Efficacy. PLoS One 10:e0118557.

21. Vallino M, Rossi M, Ottati S, Martino G, Galetto L, Marzachì C, Abbà S. 2021. Bacteriophage-host association in the phytoplasma insect vector Euscelidius variegatus. Pathogens 10.

22. Andrews S. 2010. FastQC: a quality control tool for high throughput sequence data https://doi.org/10.12688/f1000research.15931.1.

23. Bolger AM, Lohse M, Usadel B. 2014. Trimmomatic: A flexible trimmer for Illumina sequence data. Bioinformatics 30:2114–2120.

24. Bankevich A, Nurk S, Antipov D, Gurevich AA, Dvorkin M, Kulikov AS, Lesin VM, Nikolenko SI, Pham SON, Prjibelski AD, Pyshkin A V, Sirotkin A V, Vyahhi N, Tesler G, Alekseyev MAXA, Pevzner PA. 2012. SPAdes: A New Genome Assembly Algorithm and Its Applications to Single-Cell Sequencing. J Comput Biol 19:455–477.

25. Wick RR, Schultz MB, Zobel J, Holt KE. 2015. Bandage: interactive visualization of de novo genome assemblies. Bioinformatics 31:3350–3352.

26. Wick RR, Judd LM, Gorrie CL, Holt KE. 2017. Unicycler: Resolving bacterial genome assemblies from short and long sequencing reads. PLoS Comput Biol 13:1–22.

27. Wattam AR, Abraham D, Dalay O, Disz TL, Driscoll T, Gabbard JL, Gillespie JJ, Gough R, Hix D, Kenyon R, Machi D, Mao C, Nordberg EK, Olson R, Overbeek R, Pusch GD, Shukla M, Schulman J, Stevens RL, Sullivan DE, Vonstein V, Warren A, Will R, Wilson MJC, Yoo HS, Zhang C, Zhang Y, Sobral BW. 2014. PATRIC, the bacterial bioinformatics database and analysis resource. Nucleic Acid Res 42:D581–D591.

28. Mcnair K, Zhou C, Dinsdale EA, Souza B, Edwards RA. 2019. PHANOTATE: A novel approach to gene identification in phage genomes. Bioinformatics 35:4537–4542.

29. Mcnair K, Aziz RK, Pusch GD, Overbeek R, Dutilh BE, Edwards R. 2018. Phage Genome Annotation Using the RAST Pipeline, p. 231–238. In Clokie M., Kropinski A., Lavigne R. (eds) Bacteriophages. Methods in Molecular Biology. Humana Press, New York, NY.

30. Zimmermann L, Stephens A, Nam SZ, Rau D, Kübler J, Lozajic M, Gabler F, Söding J, Lupas AN, Alva V. 2018. A Completely Reimplemented MPI Bioinformatics Toolkit with a New HHpred Server at its Core. J Mol Biol 430:2237–2243.

31. Thorvaldsdóttir H, Robinson JT, Mesirov JP. 2013. Integrative Genomics Viewer (IGV): high-performance genomics data visualization and exploration. Brief Bioinform 14:178–192.

32. Coppens L, Lavigne R. 2020. SAPPHIRE: A neural network based classifier for σ70 promoter prediction in Pseudomonas. BMC Bioinformatics 21:1–7.

33. Bailey, Timothy L, Johnson J, Grant CE, Noble WS. 2015. The MEME Suite. Nucleic Acids Res 43:W39–W49.

34. Bailey TL, Gribskov M. 1998. Combining evidence using p-values: Application to sequence homology searches. Bioinformatics 14:48–54.

35. Naville M, Ghuillot-Gaudeffroy A, Marchais A, Gautheret D. 2011. ARNold: A web tool for the prediction of rho-independent transcription terminators. RNA Biol 8:11–13.

36. Mejía-Almonte C, Busby SJW, Wade JT, van Helden J, Arkin AP, Stormo GD, Eilbeck K, Palsson BO, Galagan JE, Collado-Vides J. 2020. Redefining fundamental concepts of transcription initiation in bacteria. Nat Rev Genet 21:699–714.

37. Lammens EM, Boon M, Grimon D, Briers Y, Lavigne R. 2022. SEVAtile: a standardised DNA assembly method optimised for Pseudomonas. Microb Biotechnol 15:370–386.

38. Beal J, Haddock-Angelli T, Baldwin G, Gershater M, Dwijayanti A, Storch M, de Mora K, Lizarazo M, Rettberg R. 2018. Quantification of bacterial fluorescence using independent calibrants. PLoS One 13:e0199432.

39. SAS Institute Inc. 2022. JMP®. Version 16. Cary, NC.

40. Cornelissen A, Ceyssens PJ, T’Syen J, van Praet H, Noben JP, Shaburova O V., Krylov VN, Volckaert G, Lavigne R. 2011. The T7-Related Pseudomonas putida Phage ϕ15 Displays Virion-Associated Biofilm Degradation Properties. PLoS One 6:e18597.

41. Sillankorva S, Neubauer P, Azeredo J. 2008. Isolation and characterization of a T7-like lytic phage for Pseudomonas fluorescens. BMC Biotechnol 8:80.

42. Storms ZJ, Teel MR, Mercurio K, Sauvageau D. 2020. The Virulence Index: A Metric for Quantitative Analysis of Phage Virulence. PHAGE Ther Appl Res 1:27–36.

43. Konopacki M, Grygorcewicz B, Dołęgowska B, Kordas M, Rakoczy R. 2020. PhageScore: A simple method for comparative evaluation of bacteriophages lytic activity. Biochem Eng J 161:107652.

44. Kovalyova I V., Kropinski AM. 2003. The complete genomic sequence of lytic bacteriophage gh-1 infecting Pseudomonas putida -Evidence for close relationship to the T7 group. Virology 311:305–315.

45. Weidel W, Koch G, Lohss F. 1954. Über die Zellmembran von Escherichia Coli B. Zeitschrift fur Naturforsch - Sect B J Chem Sci 9:398–406.

46. Chibeu A, Ceyssens PJ, Hertveldt K, Volckaert G, Cornelis P, Matthijs S, Lavigne R. 2009. The adsorption of Pseudomonas aeruginosa bacteriophage φKMV is dependent on expression regulation of type IV pili genes. FEMS Microbiol Lett 296:210–218.

47. Lavigne R, Lecoutere E, Wagemans J, Cenens W, Aertsen A, Schoofs L, Landuyt B, Paeshuyse J, Scheer M, Schobert M, Ceyssens PJ. 2013. A multifaceted study of Pseudomonas aeruginosa shutdown by virulent podovirus LUZ19. MBio 4.

48. Hausmann R. 1988. The T7 Group, p. 259–290. In Calendar, R (ed.), The Bacteriophages, Volume 1. Plenum Press, New York.

49. Nishimura Y, Yoshida T, Kuronishi M, Uehara H, Ogata H, Goto S. 2017. ViPTree: the viral proteomic tree server. Bioinformatics 33:2379–2380.

50. Cheetham GMT, Steitz TA. 2020. Structure of a transcribing T7 RNA polymerase initiation complex. Struct Insights into Gene Expr Protein Synth 286:301–305.

51. Cheetham GM, Steitz TA. 2000. Insights into transcription: Structure and function of single-subunit DNA-dependent RNA polymerases. Curr Opin Struct Biol 10:117–123.

52. Dunn JJ, Studier FW. 1983. Complete nucleotide sequence of bacteriophage T7 DNA and the locations of T7 genetic elements. J Mol Biol 166:477–535.

53. Lammens E-M, Nikel PI, Lavigne R. 2020. Exploring the synthetic biology potential of bacteriophages for engineering non-model bacteria. Nat Commun 11.

54. Cheng X, Zhang X, Pflugrath JW, Studier FW. 1994. The structure of bacteriophage T7 lysozyme, a zinc amidase and an inhibitor of T7 RNA polymerase. Proc Natl Acad Sci U S A 91:4034–4038.

55. Huang J, Villemain J, Padilla R, Sousa R. 1999. Mechanisms by which T7 lysozyme specifically regulates T7 RNA polymerase during different phases of transcription. J Mol Biol 293:457–475.

56. Boon M, De Zitter E, De Smet J, Wagemans J, Voet M, Pennemann FL, Schalck T, Kuznedelov K, Severinov K, Van Meervelt L, De Maeyer M, Lavigne R. 2020. “Drc”, a structurally novel ssDNA-binding transcription regulator of N4-related bacterial viruses. Nucleic Acids Res 48:445–459.

57. Stern A, Mayrose I, Penn O, Shaul S, Gophna U, Pupko T. 2010. An Evolutionary Analysis of Lateral Gene Transfer in Thymidylate Synthase Enzymes. Syst Biol 59:212–225.

58. Dwivedi B, Xue B, Lundin D, Edwards RA, Breitbart M. 2013. A bioinformatic analysis of ribonucleotide reductase genes in phage genomes and metagenomes. BMC Evol Biol 13.

59. Clokie MRJ, Mann NH. 2006. Marine cyanophages and light. Environ Microbiol 8:2074–2082.

60. Rihtman B, Bowman-Grahl S, Millard A, Corrigan RM, Clokie MRJ, Scanlan DJ. 2019. Cyanophage MazG is a pyrophosphohydrolase but unable to hydrolyse magic spot nucleotides. Environ Microbiol Rep 11:448–455.

61. Wagemans J, Blasdel BG, Van Den Bossche A, Uytterhoeven B, De Smet J, Paeshuyse J, Cenens W, Aertsen A, Uetz P, Delattre A-S, Ceyssens P-J, Lavigne R. 2014. Functional elucidation of antibacterial phage ORFans targeting Pseudomonas aeruginosa https://doi.org/10.1111/cmi.12330.

62. Summer EJ, Berry J, Tran TAT, Niu L, Struck DK, Young R. 2007. Rz/Rz1 Lysis Gene Equivalents in Phages of Gram-negative Hosts. J Mol Biol 373:1098–1112.

63. Fernandes S, São-José C. 2018. Enzymes and mechanisms employed by tailed bacteriophages to breach the bacterial cell barriers. Viruses 10:1–22.

64. Yan B, Boitano M, Clark TA, Ettwiller L. 2018. SMRT-Cappable-seq reveals complex operon variants in bacteria. Nat Commun 9.

65. Mao X, Ma Q, Liu B, Chen X, Zhang H, Xu Y. 2015. Revisiting operons: An analysis of the landscape of transcriptional units in E. coli. BMC Bioinformatics 16.

66. Lee Y, Lee N, Jeong Y, Hwang S, Kim W, Cho S, Palsson BO, Cho BK. 2019. The Transcription Unit Architecture of Streptomyces lividans TK24. Front Microbiol 10:1–13.

67. Shitrit D, Hackl T, Laurenceau R, Raho N, Carlson MCG, Sabehi G, Schwartz DA, Chisholm SW, Lindell D. 2021. Genetic engineering of marine cyanophages reveals integration but not lysogeny in T7-like cyanophages. ISME J 2021 162 16:488–499.

68. Martínez-García E, Jatsenko T, Kivisaar M, de Lorenzo V. 2015. Freeing Pseudomonas putida KT2440 of its proviral load strengthens endurance to environmental stresses. Environ Microbiol 17:76–90.

69. Bailly-Bechet M, Vergassola M, Rocha E. 2007. Causes for the intriguing presence of tRNAs in phages. Genome Res 17:1486.

70. Williams KP. 2002. Integration sites for genetic elements in prokaryotic tRNA and tmRNA genes: sublocation preference of integrase subfamilies. Nucleic Acids Res 30:866–875.

71. Brives C, Pourraz J. 2020. Phage therapy as a potential solution in the fight against AMR: obstacles and possible futures. Palgrave Commun 2020 61 6:1–11.

